# DLK-dependent biphasic reactivation of herpes simplex virus latency established in the absence of antivirals

**DOI:** 10.1101/2022.02.25.482019

**Authors:** Sara Dochnal, Husain Y. Merchant, Austin R. Schinlever, Aleksandra Babnis, Daniel P. Depledge, Angus C. Wilson, Anna R. Cliffe

## Abstract

Understanding the molecular mechanism of herpes simplex virus type 1 (HSV-1) latent infection and reactivation in neurons requires the use of model systems. However, establishing a quiescent infection in cultured neurons is problematic as infectious virus released from any productively infected neuron can superinfect the cultures. Here, we describe a new reporter virus, HSV-1 Stayput-GFP, that is defective for cell-to-cell spread and can be used to model latency and reactivation at the single neuron level. Importantly, quiescent infection of neurons with Stayput-GFP can be established without the use of a viral DNA replication inhibitor. The establishment of a quiescent state requires a longer time frame than previous models of HSV latency using DNA replication inhibitors. This results in a decreased ability of the virus to reactivate, and the use of multiple reactivation triggers is required. Using this system, we demonstrate that an initial Phase I wave of lytic gene expression occurs independently of histone demethylase activity and viral DNA replication but is dependent on the neuronal cell stress protein, DLK. Progression into the later, Phase II wave of reactivation, characterized by detectable late viral protein synthesis and a requirement for histone demethylase activity, is also observed with the Stayput-GFP model system. These data demonstrate that the two waves of viral gene expression following HSV-1 reactivation are independent of secondary infection and occur when latent infections are established in the absence of a viral DNA replication inhibitor.

**Importance:** Herpes simplex virus-1 (HSV-1) enters a latent infection in neurons and periodically reactivates. Reactivation manifests as a variety of clinical symptoms. Studying latency and reactivation *in vitro* is invaluable, allowing the molecular mechanisms behind both processes to be targeted by therapeutics that reduce the clinical consequences. Here, we describe a novel *in vitro* model system using a cell-to-cell spread defective HSV-1, known as Stayput-GFP, which allows for the study of latency and reactivation at the single neuron level. We anticipate this new model system will be an incredibly valuable tool for studying the establishment and reactivation of HSV-1 latent infection *in vitro*. Using this model, we find that initial reactivation events are dependent on cellular stress kinase DLK, but independent of histone demethylase activity and viral DNA replication. Our data therefore demonstrate the essential role of DLK in mediating a wave of lytic gene expression unique to reactivation.

## Introduction

Herpes simplex virus 1 (HSV-1) is a globally prevalent pathogen with the capacity to infect both sensory and autonomic neurons (1–4) . Following neuronal infection, HSV-1 can enter a lytic replication cycle, establish a lifelong latent infection, or potentially undergo a limited amount of lytic gene expression even prior to latency establishment (5–13). While latency is largely asymptomatic, periodic reactivation of the virus can result in cutaneous lesions, keratitis, and encephalitis. Epidemiological studies have also linked HSV infection with an increased risk of developing late onset Alzheimer’s disease (14–22).

The regulated expression of viral lytic transcripts has been well characterized following lytic infection (23–25). The HSV-1 genome enters the host cell epigenetically naked (26–28) but becomes chromatinized by histones bearing transcriptionally permissive histone modifications (29–38). Viral gene expression is initiated from the genome in response to viral trans-activator and tegument protein VP16, which forms a complex with cellular factors involved in transcriptional activation, including general transcription factors, ATP-dependent chromatin remodelers, and histone modifying enzymes to promote expression of immediate-early (IE) genes (32, 39–43). Synthesis of the IE proteins is required for early (E) mRNA transcription. Products of the early viral genes enable viral DNA replication. Viral genome synthesis is a pre-requisite for true-late (TL) mRNA transcription, likely due to a shift in genome accessibility and increased binding of host transcriptional machinery (44). In contrast to productive infection, during HSV-1 latency, viral lytic mRNAs are largely transcriptionally repressed and promoters assemble into silent heterochromatin marked by the tri-methylation of histone H3 lysine 27 (H3K27me3) and di/tri-methylation of histone H3 on lysine 9 (H3K9me2/3) (45–50). The initiation of viral gene expression during reactivation is induced from a heterochromatin-associated viral genome and occurs in the absence of viral activators like VP16. Latent HSV-1 therefore relies on host factors to act on the epigenetically silent viral genome and induce lytic gene expression.

The mechanisms that regulate entry into lytic gene expression to permit reactivation remain elusive. Using primary neuronal models of HSV-1 latent infection, reactivation has been found to progress in a two-step or bi-phasic manner. Phase I is characterized as a synchronous wave of lytic viral transcripts, occurring approximately 20 hours post-stimulus (25, 51–54). There is evidence that this initial induction of lytic gene expression is not dependent on the lytic trans-activator VP16, or viral protein synthesis (25, 55). Instead, cellular factors, including the stress kinases dual leucine zipper kinase (DLK) and c-Jun N terminal kinase (JNK), are required for Phase I entry (52, 54). Importantly, viral gene expression occurs despite the persistence of heterochromatin on viral promoters. Instead, JNK-dependent histone phosphorylation of histone H3S10 results in a methyl/phospho switch, which can permit gene expression even while the repressive H3K9me3 histone modification is maintained (52). The second wave of viral lytic gene expression, Phase II, occurs approximately 48 hours post-stimulus. Phase II reactivation is characterized by the full transcriptional viral cascade, including viral DNA replication, and ultimately infectious virus production (25). In contrast to Phase I, viral protein synthesis, heterochromatin removal, VP16-mediated transactivation, and viral DNA replication are required for Phase II (25, 52, 54).

*In vitro* systems of HSV-1 latency are required to study the mechanisms of latency establishment and reactivation that cannot be easily studied *in vivo* (56–59). *In vitro* models more readily enable to functional studies, and immune system components can be included to understand how the host immune response impacts latency and reactivation (60, 61). However, there are complications involved in establishing latency *in vitro*. As also observed in animal models, only a sub-population of infected neurons enter a latent state whereas other neurons become lytically infected (11, 62–64). This leads to superinfection of the cultures and an inability to establish a latent infection. Therefore, many existing *in vitro* models have used viral DNA replication inhibitors, predominantly acyclovir (ACV), to establish latency *in vitro* (52, 65–70). ACV is proposed to inhibit viral DNA replication by incorporating into actively replicating viral genomes in lytic cells, although there is also evidence from ACV-resistant strains that ACV can also inhibit the viral DNA polymerase (71–73). There are some caveats associated with ACV use as the process of DNA replication and subsequent late gene expression cannot be studied in lytic neurons during latency establishment. Moreover, the fate of any genomes that incorporate ACV is unknown. Therefore, new model systems are required in which latency establishment can be tracked without the need for viral DNA replication inhibitors. Such a system can also be used to determine whether the use of ACV in the cultures alters mechanisms of lytic gene expression during reactivation. Here we describe the use of a novel HSV-1 reporter virus Stayput-GFP, which provides a powerful new methodology to investigate the establishment and reactivation from latent infection at the single neuron level without the need for DNA replication inhibitors.

## Results

### Construction of a gH-null US11-GFP HSV-1

To construct a recombinant HSV-1 that is deficient in cell-to-cell spread and can also be used to visualize cells containing detectable viral late protein, Us11 tagged with GFP (74) was inserted into an existing glycoprotein H (gH)-null virus, SCgHZ (75) (Fig. 1A). gH is essential for HSV-1 cell entry as it mediates fusion between the virus envelope and host cell membrane (76, 77). The GFP-tagged true-late protein is a useful indicator of both lytic infection and reactivation, and Us11-GFP wild-type virus has been used as such in several HSV-1 latency systems (51, 54, 58, 65, 78, 79). The tagged virus has previously been reported to express the full complement of HSV-1 genes, as wild-type Patton strain (74).

**FIG 1:**
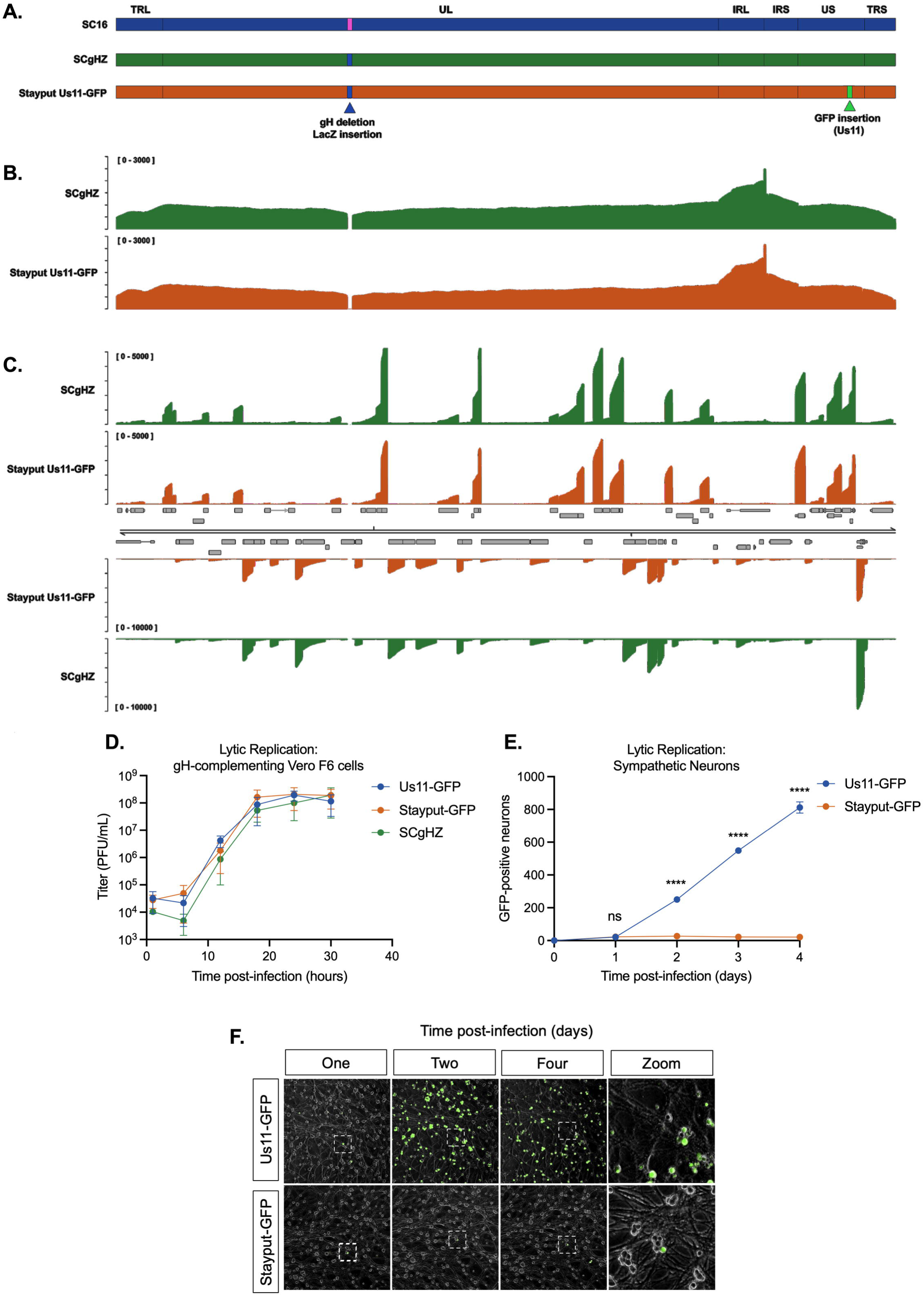
Stayput-GFP replicates as wild-type but is unable to spread. (A) Schematic overview of HSV-1 strain SC16, the gH-deletion mutant SCgHZ, and Stayput-GFP. The gH deletion / LacZ insertion and Us11-GFP insertion sites are shown by blue and green triangles, respectively. (B-C) Coverage plots derived from (B) nanopore gDNA sequencing and (C) nanopore direct RNA sequencing of SCgHZ and Stayput-GFP. Sequence read data were aligned against the SC16 reference genome and demonstrate a drop in coverage at the gH locus. (D) Vero-F6 cells were infected with Stayput-GFP, SCgHZ, or Us11-GFP at an MOI of 5. Infectious virus was collected over time and titrated on Vero-F6 cells (n=3 biological replicates). (E-F) Neonatal sympathetic neurons were infected at an MOI of 0.5 PFU/cell with Stayput-GFP or Us11-GFP in the absence of DNA replication inhibitors. Us11-GFP-positive neurons were counted over time (n=3 biological replicates). Shapiro-Wilk normality test. Unpaired student’s t test between Us11-GFP and Stayput-GFP. ****p<0.0001. The means and SEMs are shown.

We first verified that the resulting virus (named Stayput-GFP) was deficient in cell-to-cell spread in non-neuronal and neuronal cells but otherwise undergoes gene expression and replication as a wild-type virus. The genome sequence and transcriptome of Stayput-GFP was validated by nanopore gDNA and direct RNA sequencing, respectively (Fig. 1B-C). As determined through plaque assay, the ability to produce infectious virus was perturbed in the gH-deletion strains but was rescued using previously constructed gH-complementing cell line, Vero-F6 (Fig. 1D, Supplemental Fig. 1A). Importantly, replication in this cell line was indistinguishable from the parent SCgHZ and Us11-GFP Patton strain, which we chose for comparison as this virus has previously been used for latent infection studies in primary neurons.

To demonstrate cell-to-cell spread deficiency in neurons, we quantified GFP-positive neurons over time. Infection of murine sympathetic neurons at an MOI of 0.5 PFU/cell resulted in detectable GFP-positive neurons, which remained constant after 24 hours post-infection (Fig. 1E), while the surrounding GFP-negative neurons remained GFP-negative (Fig. 1F). This contrasts with the wild-type US11-GFP condition where the number of lytic-infected neurons increased substantially over time. At 6 hours post-infection, viral protein ICP4-positive neurons were indistinguishable between SCgHZ, Stayput-GFP, or wild-type Us11-GFP infected neurons (Supplemental Fig. 1B). Therefore, Stayput-GFP neuronal infection was equivalent to Us11-GFP upon initial infection but failed to spread within the neuronal culture.

### Reactivation of Stayput-GFP in a primary neuronal model

To confirm that Stayput-GFP undergoes reactivation in a manner comparable to the backbone SCgHZ, we infected mouse primary sympathetic neurons in the presence of viral DNA replication inhibitor ACV (52, 65–68). ACV prevents the production of infectious virus, and thus superinfection of the cultures. Following infection, ACV was removed, and reactivation was triggered adding a PI3-kinase inhibitor LY294002 (Fig. 2), which mimics loss of a branch of the NGF-signaling pathway in neurons (80) and has previously been found to induce reactivation (52, 61, 65, 79). Following the addition of the reactivation stimulus, viral gene expression increased uniformly between Stayput-GFP, SCgHZ, and wild-type Us11-GFP at 20 hours post-stimulus (Fig. 2C). Viral gene expression continued to increase from 20 to 48 hours for all three viruses, and GFP-positive neurons were visible for both GFP-tagged viruses by 48 hours (Fig. 2B, 2D). This indicates that even in the absence of cell-to-cell spread, reactivation progressed over a 48-hour period, initiating with viral mRNA production and later detection of viral late protein.

**FIG 2:**
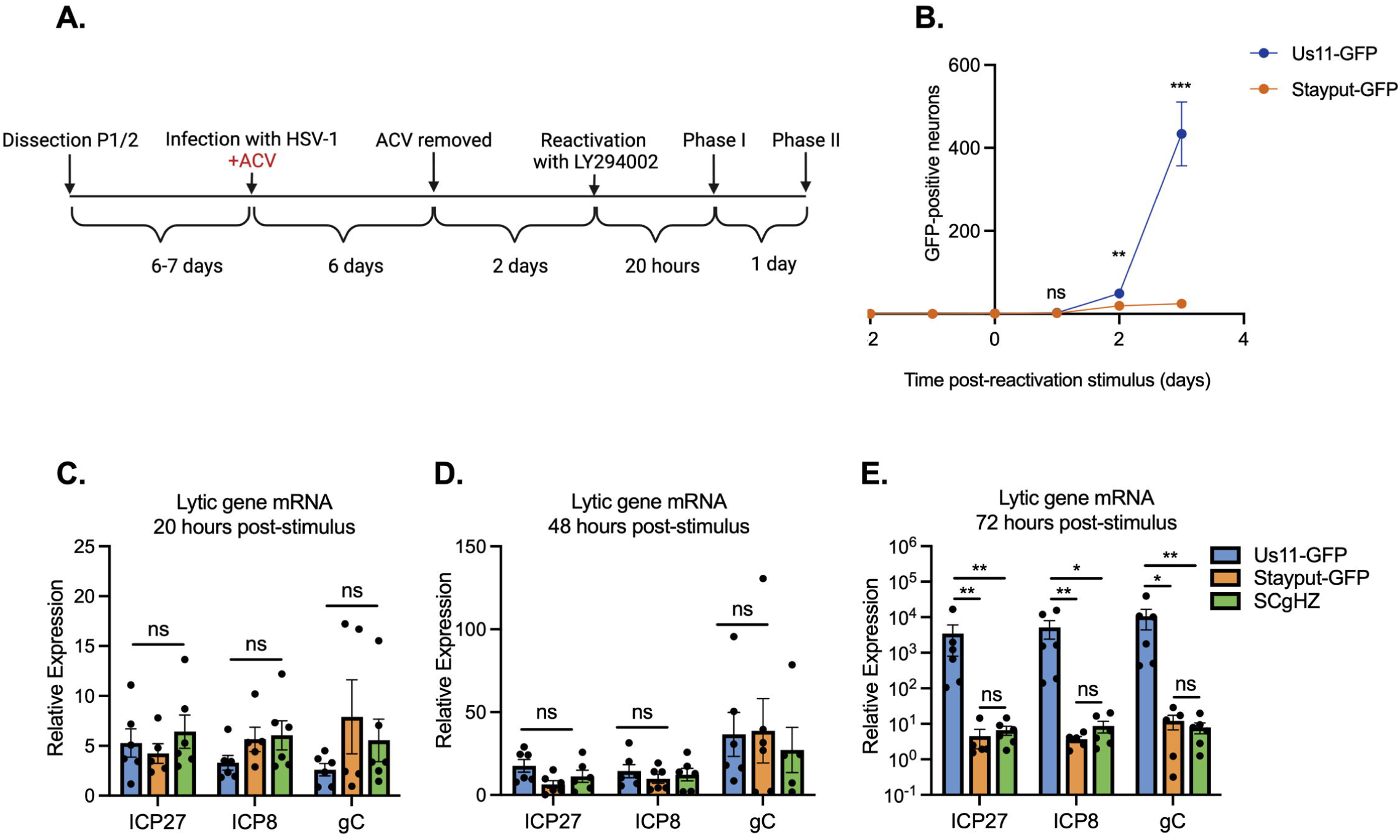
Stayput-GFP in a latency and reactivation model using ACV to promote latency establishment. (A) The latency and reactivation model scheme. Neonatal sympathetic neurons were infected with Stayput-GFP, parent virus SCgHZ, or wild-type Us11-GFP at an MOI of 7.5 PFU/cell in the presence of ACV (50 μM). 6 days later, ACV was removed, and 2 days later, cultures were reactivated with LY294002 (20 μM). The numbers of GFP positive neurons in a single well (containing approximately 5,000 neurons) for Stayput-GFP and wild-type Us11-GFP were counted over time (B). Viral gene expression also was quantified by RT-qPCR for immediate early (*ICP27*), early (*ICP8*), and late (*gC*) genes at 20 hours (C), 48 hours (D), and 72 hours (E) post-stimulus. Relative expression to un-reactivated samples and cellular control (mGAPDH). n=6 biological replicates from 3 litters. Normality determined by Kolmogorov-Smirnov test (B-E). Mann-Whitney (B) or Kruskal-Wallis with comparison of means (C-E). *p<0.05, **p<0.01, ***p<0.001. The means and SEMs are represented. Individual biological replicates are indicating in C-E.

By 72 hours-post stimulus, GFP and viral gene transcription were significantly up-regulated in wild-type Us11-GFP in comparison to Stayput-GFP or SCgHZ, suggesting that at this time-point, the readout of reactivation for wild-type Us11-GFP is confounded by cell-to-cell spread (Fig. 2B, 2E). We are therefore unable to differentiate between genuine reactivation and downstream cell-to-cell spread using a wild-type virus. Previous attempts to reduce cell-to-cell spread include using pooled human gamma globulin or the viral DNA packaging inhibitor WAY-150138 (65, 81–83). However, using Stayput-GFP offers a built-in mechanism to prevent cell-to-cell spread during reactivation and permit quantification of the progression to reactivation at the single neuron level.

### Stayput-GFP can be used to create a quiescence model of neuronal infection in the absence of viral DNA replication inhibitors

When wild-type virus is used for neuronal infection, superinfection of the cultures occurs (Fig. 1E), and a latent infection cannot be established. By infecting with Stayput-GFP, we posited that we could create a model of latency establishment without the use of DNA replication inhibitors. Following the infection of neonatal sympathetic neurons at an MOI of 7.5 PFU/cell, the number of GFP-positive neurons emerged by 1-day post-infection, increased until 3 days post-infection, and then decreased until reaching zero by 30 days post-infection (Fig. 3A). The length of time required for Us11-GFP to be lost from the cultures was surprising and may result from the previously characterized restricted cell death in neurons (84), or a gradual shut-off in protein synthesis in a sub-population of neurons that survive even Late gene expression. Single-cell tracking demonstrates that most neurons that become GFP-positive also end up staining positive for cell death marker SYTOX^TM^ Orange (Fig. 3B-C). All classes of lytic viral gene expression emerged by 1-day post-infection and then decreased over the span of 30-days post-infection (Fig. 3D-F). In contrast to the lytic transcripts, LAT expression was maintained over the infection scheme of 30-40 days and was approximately 400-fold higher than lytic transcripts from 30 days onwards (Fig. 3G). This indicated that LAT-positive neurons persisted over this period, likely reflecting entry into quiescence. In agreement with this hypothesis, at 40 days post-infection there are approximately 200 viral DNA copies per cell, demonstrating that viral genomes persist (Fig. 3H). Together, these data show that infection of sympathetic neurons with a cell-to-cell defective virus results in a remaining population of neurons containing viral genomes and the LAT transcript. Notably, this mimics events following *in vivo* infection of mice.

**FIG 3:**
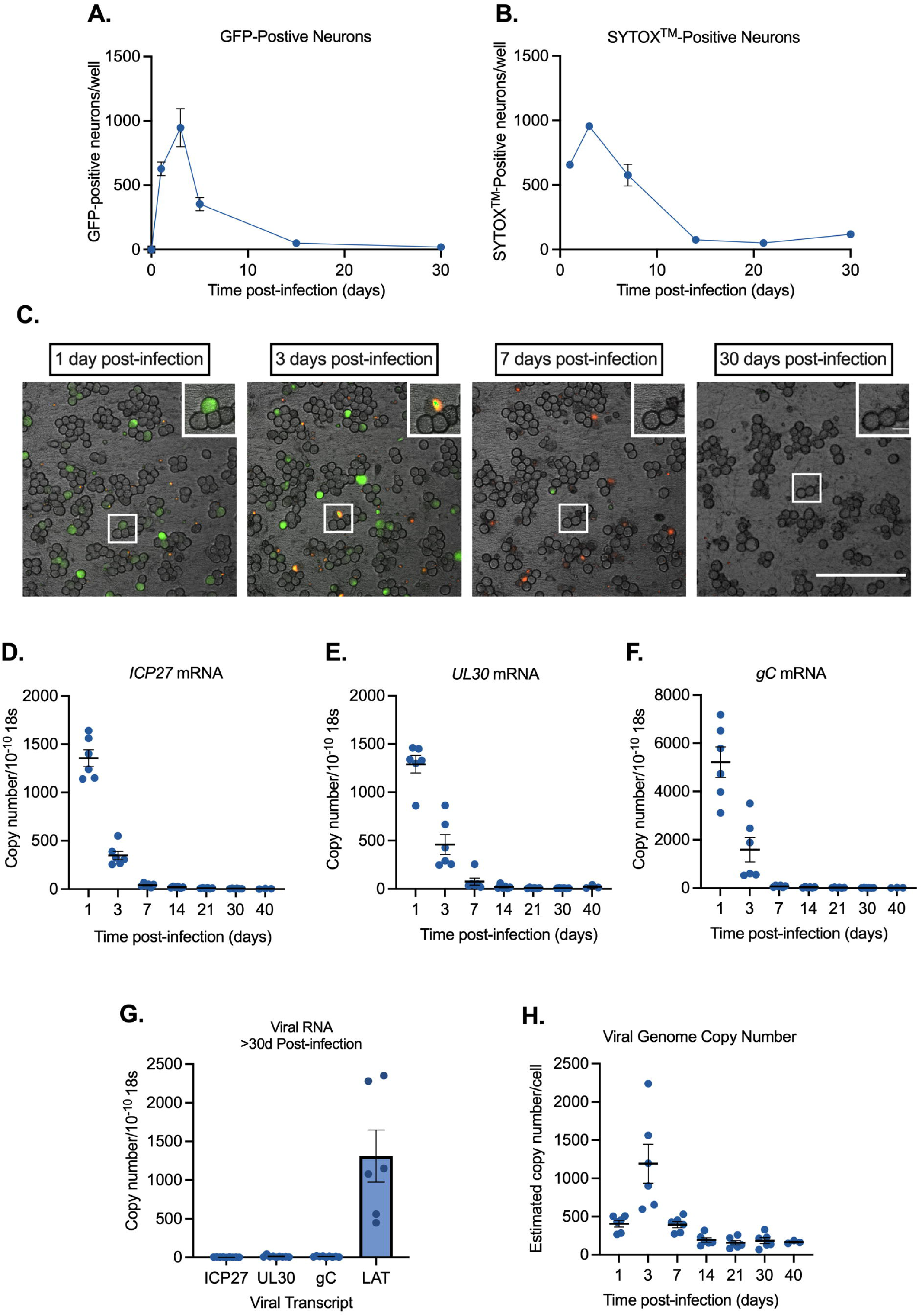
Stayput-GFP can be used to create a quiescence model in the absence of viral DNA replication inhibitors in neonatal sympathetic neurons. Neonatal sympathetic neurons were infected with Stayput-GFP at an MOI of 7.5 PFU/cell and the numbers of Us11-GFP-positive neurons were quantified. n=9 biological replicates from 3 litters (A). SYTOX^TM^ Orange-positive neurons were also quantified over time (n=3) (B). Following infection, the same field of view was imaged to track GFP and SYTOX^TM^ Orange (250 μm scale bar for FOV, 25 μm scale bar for zoom) over time (C). Lytic (D-F) and latent (G) viral transcripts (n=6) were quantified up to 40 days post-infection. Viral DNA load (n=6) (H) was also quantified up to 40 days post-infection. Individual biological replicates along with the means and SEMs are shown.

We were also interested to determine whether a similar quiescent infection could be established in adult sensory neurons, as these are also a commonly used model of HSV-1 infection *in vitro* (62). Following infection with Stayput-GFP in primary trigeminal ganglia (TG) neurons, GFP also increased by 1-day post-infection and then was lost over time (Supplemental Fig. 2). GFP was repeatedly lost within 15 days, a shorter period than that which is observed in neonatal sympathetic neurons. Therefore, the Stayput-GFP virus can be used to establish a quiescent infection of both sympathetic and sensory neurons *in vitro* without the use of viral DNA replication inhibitors.

### The ability of HSV-1 to undergo reactivation decreases with length of time infected

The presence of viral genomes and LAT transcripts suggested that HSV-1 had established a quiescent infection 30 days post-infection. Therefore, we hypothesized that some genomes enter latency, which is defined by an ability to reactivate in response to a stimulus. We thus attempted to reactivate cultures with LY294002 (20 μM). However, we were unable to detect an increase in GFP-positive neurons after the addition of the trigger (Fig. 4A). We were also unable to detect a change in immediate early, early, or late transcripts (data not shown). This was unexpected as LY294002 has repeatedly been shown to elicit robust reactivation *in vitro* and was able to induce Stayput-GFP in a model using ACV to promote latency establishment (Fig. 2). Therefore, we sought to determine whether the inability to induce reactivation was due to the lack of ACV or the more prolonged time between initial infection and the addition of the reactivation stimulus. We infected neonatal cultures in the presence of ACV and reactivated over increasing lengths of time. ACV was removed from all cultures 6 days post-infection. We found that the number of GFP-positive neurons following addition of LY294002 decreased as the length of time infected increased (Fig. 4B). This was not likely due to a loss of viral genomes, as viral genome copy number and LAT expression remained constant over this period (Fig. 3G-H) and therefore instead reflected a more repressed viral genome unable to undergo reactivation upon PI3-kinase inhibition.

**FIG 4:**
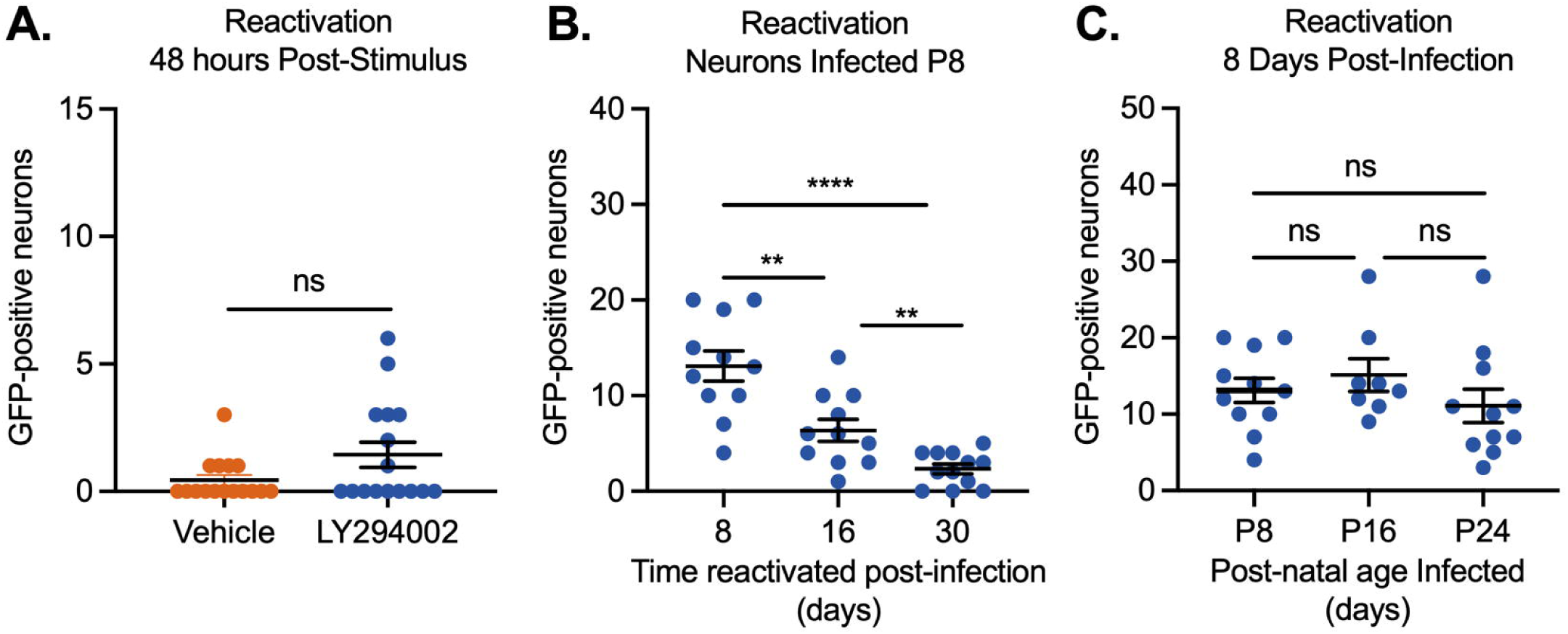
Reactivation decreases with length of time infected. Sympathetic neurons were infected with Stayput-GFP at an MOI of 7.5 PFU/cell and were treated with LY294002 when GFP-positive neurons were no longer detected (approximately 30 days post-infection). GFP-positive neurons were quantified over time; peak GFP (48 hours post-stimulus) is represented (A). Neonatal SCGs were infected at age postnatal day 8 (P8) with Stayput-GFP in the presence of ACV. ACV was removed 6 days post-infection and reactivation was triggered at the indicated times post-infection (B). Neonatal SCGs were infected as above after different lengths of time *in vitro*, representing indicated postnatal ages, and reactivated 8 days post-infection with LY294002 (C). n=12 biological replicates from 3 litters. Normality determined by Kolmogorov-Smirnov test. Unpaired student’s t-test (B) or Mann-Whitney (A, C) based on normality of data. **p<0.01, ***p<0.001, ****p<0.0001. Individual biological replicates along with the means and SEMs are shown.

Neurons are known to undergo intrinsic maturation, even in culture (85, 86). Therefore, the increased age of the neuron could have also impacted the ability of HSV-1 to undergo reactivation. Hence, we investigated how reactivation changed with increased neuronal maturation (Fig. 4C). We infected cultured neurons with Stayput-GFP at an MOI of 7.5 at the postnatal (P) ages of P8, P16, and P24 and then reactivated 8 days later. These postnatal ages of infection were chosen to reactivate at the same ages of reactivation in Fig 4B. Importantly, we did not detect a decrease in reactivation output as the age of the neuron increased. Together, these data indicate that the decreased ability of Stayput-GFP to reactivate in a model that did not require ACV to establish quiescence was due to the longer time frame of infection and not associated with a lack of ACV in the cultures or increased age of the neuron.

### Viral gene expression can be induced following long-term quiescent infection when multiple triggers are combined

We next sought to determine whether other known stimuli of HSV-1 reactivation could induce Us11-GFP expression, indicative of entry into reactivation. We attempted a number of triggers including forskolin (87–90) and heat-shock/hyperthermia (91–98), which are both known inducers of HSV-1 reactivation. Alone these stimuli did not induce Us11-GFP expression or viral lytic mRNA induction (data not shown). However, when heat-shock (43°C for 3h), in addition to forskolin (30 μM) and LY294002 (both pulsed for 20 hours) were combined, Us11-GFP positive neurons were detected at 48 hours post-stimulus, indicating progression to reactivation (Fig. 5A). Superinfection was also used as this can induce rapid and robust reactivation, likely resulting in delivery of viral tegument proteins. In comparison to superinfection, the combined heat-shock/forskolin/LY294002 trigger resulted in reduced entry into reactivation/Us11-GFP expression, indicating that only a sub-population of neurons undergo reactivation with this combined trigger, which is consistent with previous studies investigating the mechanism of HSV-1 reactivation both *in vivo* and *in vitro* (25, 66, 99, 100). Importantly, GFP-positive neurons at 48 hours post-stimulus co-stained with viral immediate early protein ICP4, early protein ICP8, as well as late capsid protein ICP5 (Supplemental Fig 3). ICP8 staining appeared to form replication compartments, suggesting viral DNA replication occurs at this time. ICP5 capsid staining was also detectable in GFP-positive axons.

**FIG 5:**
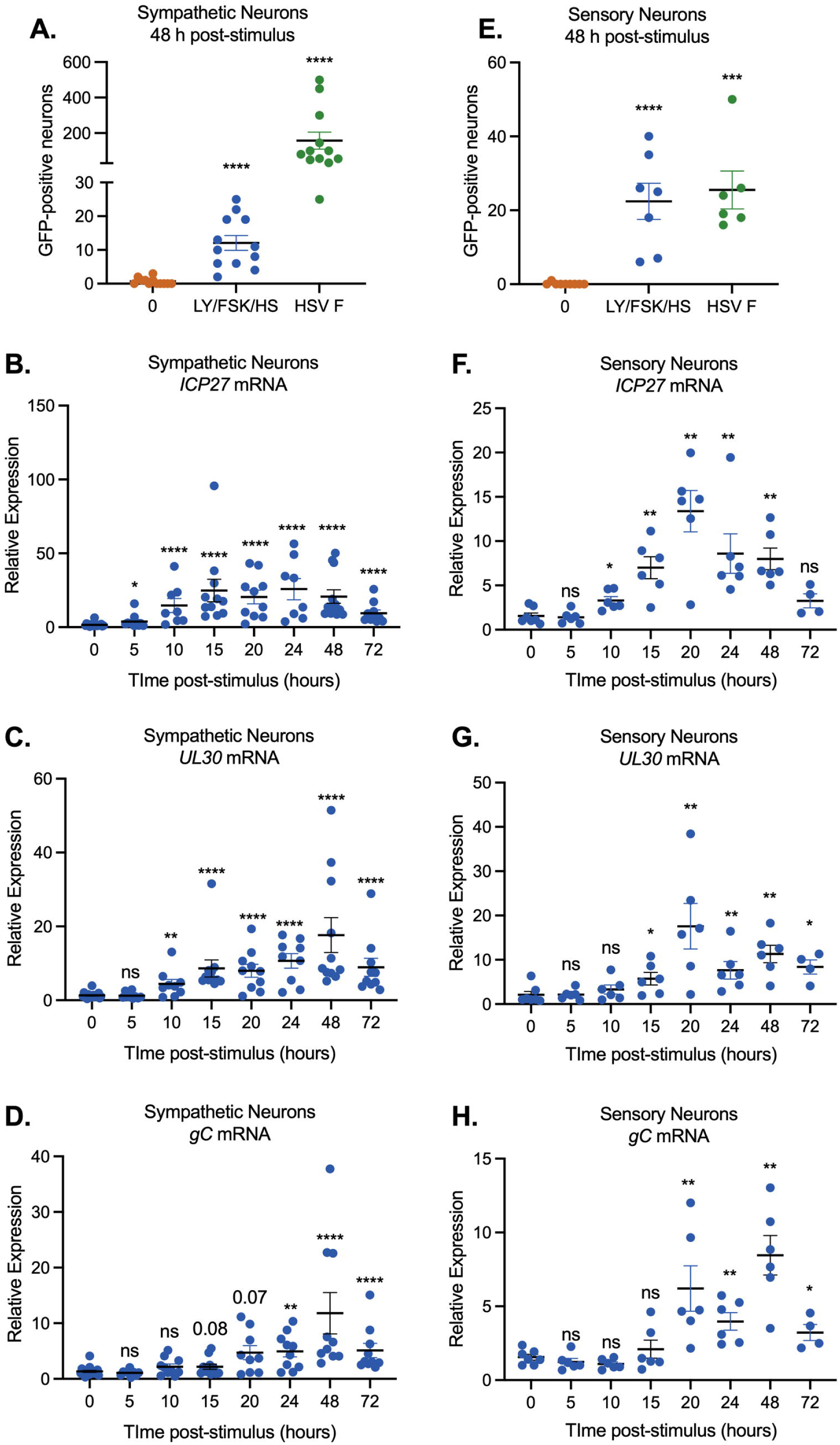
Viral gene expression can be restarted following latency establishment. Neonatal sympathetic neurons or adult sensory neurons were infected at an MOI of 7.5 PFU/cell with Stayput-GFP in the absence of viral DNA replication inhibitors. Following the loss of GFP, signaling quiescence of the culture, wells were reactivated with a variety of triggers, including combinations of LY-294002 (20 μM), forskolin (60 μM), and heat shock (43°C for 3 hours), as well as a superinfection with untagged F strain at an MOI of 10 PFU/cell. GFP was quantified over time, and the peak GFP, at 48 hours post-stimulus, is depicted. N=12 biological replicates (A, E). Immediate early (B, F), early (C, G) and late (D, H) viral transcripts were investigated over time following the stimulus. Neonatal sympathetic neurons n=9 biological replicates from 3 litters. Mann-Whitney against 0 hours (A-D). Adult sensory neurons n=9 biological replicates (E) or n=6 replicates (F, G, H). Normality determined by Kolmogorov-Smirnov test. Mann-Whitney against 0 hours. *p<0.05, **p<0.01, ***p<0.001, ****p<0.0001. Individual biological replicates along with the means and SEMs are shown.

We went on to investigate whether viral mRNA expression was induced prior to the detection of Us11-GFP positive neurons. Using the triple stimulus, we detected an increase in viral lytic transcripts as early as 10 hours post-stimulus, which continued to increase, peaking at 48 hours post-stimulus (Fig. 5B-D, Supplemental Fig. 4). Detection of immediate-early and early transcripts was slightly more robust than late transcripts, although all late transcripts were significantly induced by 20 hours post-stimulus (Fig. 5D, Supplemental Fig. 4B-E).

We also confirmed that the triple combinatorial stimulus elicits robust GFP re-emergence in adult TGs (Fig. 5E). Similar to what we observed in the sympathetic neurons, Us11-GFP positive neurons were detected by 48 hours post-stimulus. Intriguingly, superinfection only induced GFP expression to equivalent levels as the triple stimuli, which may be reflective of the repressive nature of sub-population of mature sensory neurons to lytic replication and reactivation (62, 63). Together, these data indicate that *in vitro* models of latency and reactivation can be established in sympathetic and sensory neurons in the absence of ACV and reactivation can be induced using a combination of triggers. In both models, a wave of lytic mRNA expression was detected prior to the appearance of Us11-GFP positive neurons (Fig. 5 F-H, Supplemental Figure 5).

### Neurons infected with Stayput-GFP undergo a DLK-dependent Phase I of reactivation

Phase I reactivation has largely been investigated using *in vitro* models in which ACV has been used to promote latency establishment. In addition, Phase I has been found to occur with the single triggers of forskolin or LY294002 (25, 52, 54). The requirement of multiple triggers for reactivation suggested that multiple cell-signaling pathways converged to have a synergistic effect and induce reactivation in this more repressive model. Therefore, we were interested to determine whether the characteristics of Phase I reactivation occurred in the model of quiescent infection established in the absence of ACV and using the more robust trigger to induce reactivation. Potential Phase I viral transcription was investigated at 12.5 hours post-stimulus by RT-qPCR and Phase II was investigated when Us11-GFP positive neurons could be detected (48 hours post-stimulus). A characteristic of Phase I expression is the requirement of the stress kinase dual leucine zipper kinase (DLK) (52, 54). Therefore, using the DLK-specific inhibitor GNE3511 4 µM (101), we investigated the effect of DLK on reactivation in our system. We found up-regulation of immediate early/early transcripts 12.5 hours post-stimulus was eliminated with the addition of the DLK inhibitor (Fig. 6A-C). Further, full reactivation, demonstrated by peak GFP expression at 48 hours, was also reduced to baseline levels upon the addition of the DLK inhibitor (Fig. 6D). Therefore, using our new model of HSV-1 latency our data demonstrate that the initiation of viral lytic gene expression is dependent on DLK activity.

**FIG 6:**
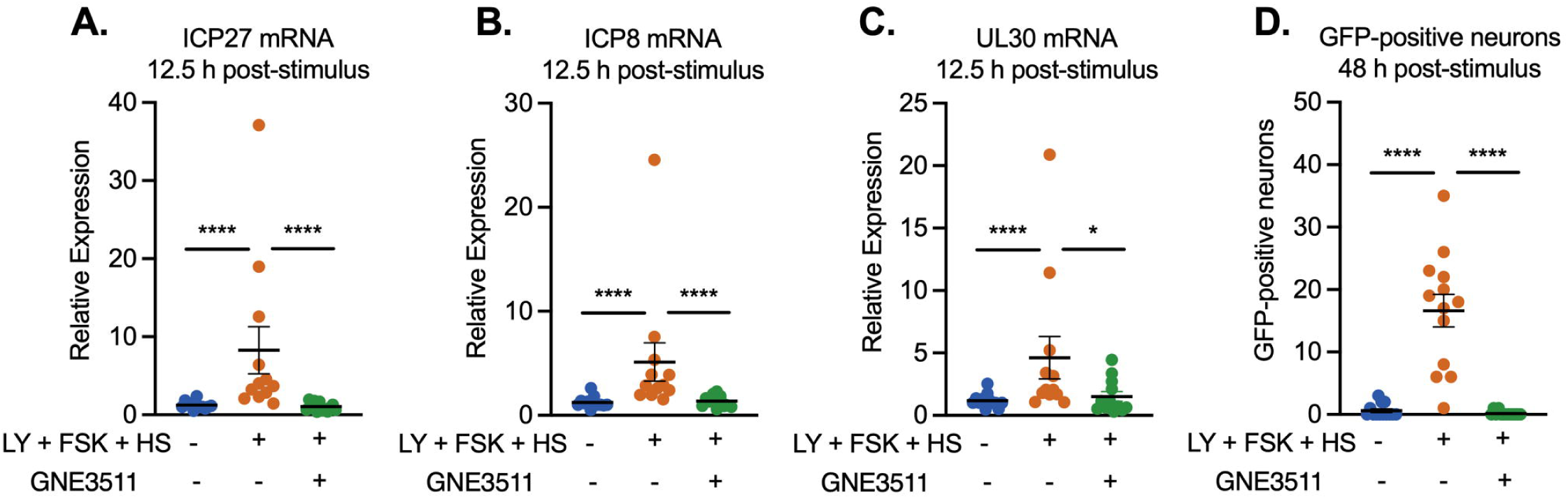
Reactivation is dependent on DLK. Cultures were infected with Stayput-GFP at an MOI of 7.5 PFU/cell in the absence of ACV. Following loss of GFP, cultures were reactivated with a combination of LY294002, forskolin, and heat shock in the presence of DLK inhibitor GNE-3511 (4 µM). Immediate early (*ICP27*) and early (*ICP8/UL30*) viral genes (A-C) were investigated at 12.5 hours-post stimulus, and GFP was counted over time (D). Peak GFP, consistently around 48 hours post-stimulus, is presented. n=9 biological replicates from 3 litters. Normality determined by Kolmogorov-Smirnov. Mann-Whitney test. (*p<0.05, **p<0.01, ***p<0.001, ****p<0.0001). The mean and SEM are shown.

In addition to the dependence on DLK activity, a further characteristic of Phase I gene expression is the induction of viral mRNA transcripts independently of the activity of histone de-methylase enzymes required for full reactivation. Therefore, we also triggered reactivation in the presence of de-methylase inhibitors. GSK-J4 is known to specifically inhibit the H3K27 histone demethylases UTX and JMJD3 (102) and has previously been found to inhibit HSV-1 reactivation but not Phase I gene expression (52, 54). OG-LOO2 is an LSD1 specific inhibitor. LSD1 has previously been shown to be involved in removal of H3K9me2 from HSV-1 genomes and its activity is required for full reactivation but not Phase I gene expression (41, 52, 54). Full reactivation was reduced in the presence of OG-LOO2 or GSK-J4 as demonstrated by GFP-positive neurons at 48 hours (Fig. 7C). However, the initial expression of lytic transcripts at 12.5 hours post-stimulation was not inhibited by either OG-LOO2 or GSK-J4 (Fig. 7A-B). Therefore, the initiation of gene expression in a manner that is independent of histone demethylase activity occurs when reactivation is induced by a triple stimulus and in a primary neuronal model in which latency was established without ACV.

**Figure 7:**
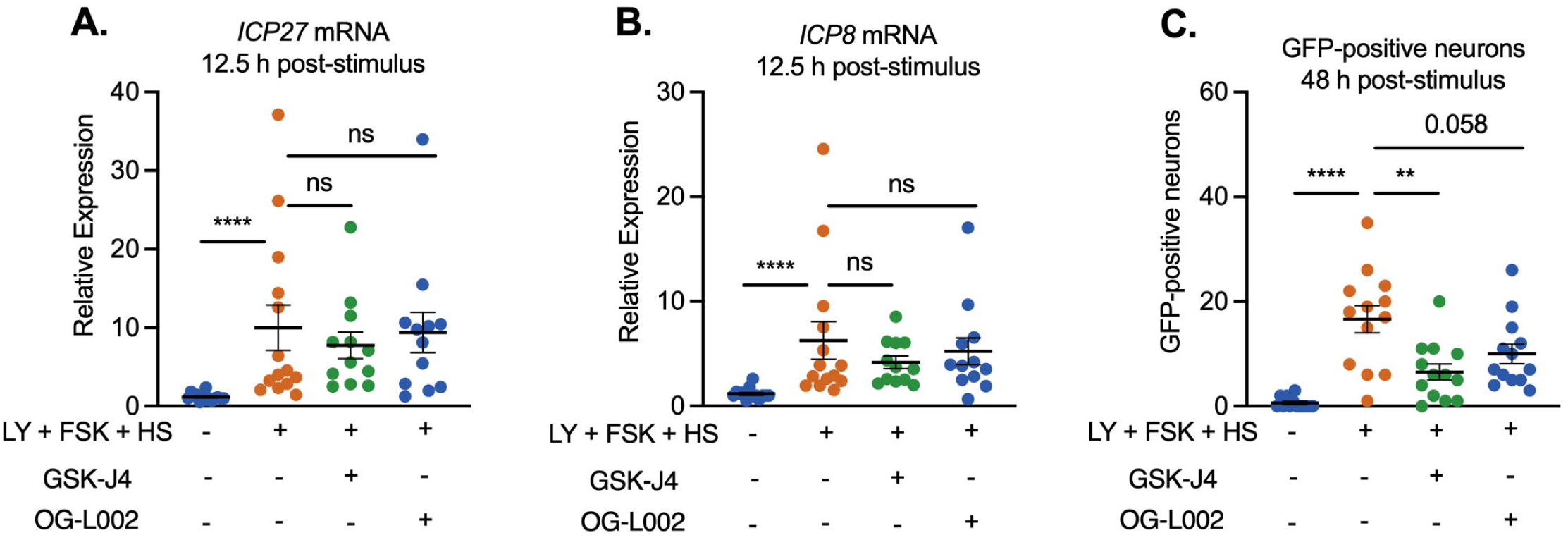
The early phase of lytic gene expression following a reactivation stimulus is independent of demethylase activity. Cultures were infected with Stayput-GFP at an MOI of 7.5 PFU/cell. Following loss of GFP, cultures were reactivated with a combination of LY-294002, forskolin, and heat shock in the presence of H3K27 demethylase inhibitor GSK-J4 (2 μM) or H3K9 demethylase inhibitor OG-L002 (20 μM). Immediate early (*ICP27*) and early (*ICP8/UL30*) viral genes (A-C) were investigated at 12.5 hours-post stimulus and GFP was counted over time (D). Peak GFP is presented. n=3 biological replicates from 3 litters. Normality determined by Kolmogorov-Smirnov. Mann-Whitney. *p<0.05, **p<0.01, ***p<0.001, ****p<0.0001. The mean and SEM are shown.

During Phase I, there is no detectable replication of viral genomes even though true late gene expression occurs (52, 54). In addition, using ACV models to establish latency, late gene expression has previously been found to occur to equivalent levels when viral DNA replication is inhibited during reactivation. Although we did observe late gene expression during the initial period of lytic gene induction (12.5-20 hours), the increase was less robust than IE and E genes and appeared slightly delayed, not reaching significance for all analyzed late transcripts until 24 hours post-stimulus (Fig. 5D, Supplemental Fig. 3). Therefore, we investigated whether this late gene induction was dependent on viral DNA replication by reactivating in the presence of ACV. The addition of ACV inhibited entry in full reactivation, demonstrated by Us11-GFP positive neurons at 48 hours post-stimulus, indicating that ACV was capable of blocking robust late gene expression at this late time-point (Fig. 8D). However, the addition of ACV did not inhibit the induction of lytic transcripts at 22 hours post-stimulus. Importantly, we included multiple true Late genes in this analysis and all were induced to equivalent levels in the presence and absence of ACV (Fig. 8A-C, Supplemental Fig 6). Therefore, the initial expression of late genes following a reactivation stimulus is independent of viral DNA replication. In summary, using a model system in which a quiescent infection is established without the need for ACV, all the previous characteristics of Phase I gene expression (dependence on DLK and independence of histone demethylase activity and viral DNA replication) were still observed.

**Figure 8:**
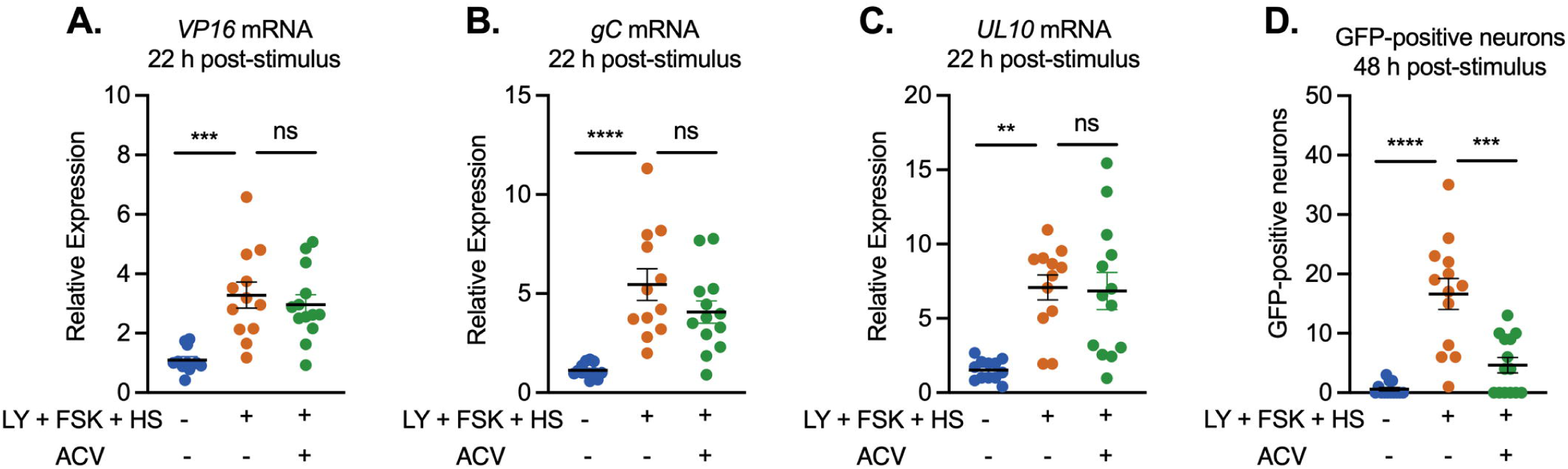
Differential dependence on viral DNA replication between Phase I and II reactivation. Cultures were infected with Stayput-GFP at an MOI of 7.5 PFU/cell in the absence of ACV. Following loss of GFP, cultures were reactivated with a combination of LY-294002, forskolin, and heat shock in the presence of ACV (50 μM). Late (*VP16, gC, UL10)*genes (A-C) were investigated at 22 hours-post stimuli. GFP was counted over time and peak GFP is presented (D). n=12 biological replicates, normality determined by Kolmogorov-Smirnov. Mann-Whitney. *p<0.05, **p<0.01, ***p<0.001, ****p<0.0001. The mean and SEM are shown.

## Discussion

We envision multiple uses for the Stayput-GFP virus model developed here for investigating HSV-1 neuronal infection *in vitro*. The Stayput-GFP virus is advantageous in models that otherwise use DNA replication inhibitors to promote latency establishment because it allows for the separation of initial viral gene expression/protein synthesis events and readouts from events that result from cell-to-cell spread. In addition, even in systems where ACV is used, there can be low levels of lytic replication or spontaneous reactivation after removal of ACV from cultures. The use of Stayput-GFP helps limit the confounding effects of spontaneous reactivation events by inhibiting subsequent cell-to-cell spread, while at the same time identifying neurons that escape quiescence. Further, the GFP tag serves as an imaging indicator in real time of when *de novo* lytic infection is resolved and latency is considered established. We are also able to track viral DNA replication and downstream late viral transcription and protein synthesis during the latency establishment process *in vitro*. The fate of lytic neurons, whether they undergo cell death or turn off gene expression programs and enter the latency pool, can also be investigated by tracking GFP and cell death at a single-cell level.

There are limitations to our system. Although there is some discrepancy in what defines reactivation (103), it is ultimately defined by the production of infectious virus. Due to the nature of the gH-deletion virus, *de novo* virus is by design non-infectious and we are unable to demonstrate reactivation in its strictest definition. That said, we can readily demonstrate the re-emergence of all classes of viral gene transcripts, synthesis of viral capsid protein and replication compartment formation.

An intriguing finding in our study is that reactivation output decreases as length of infection increases. A potential explanation is that the viral genome becomes increasingly chromatinized over time, leading to a more repressive phenotype. In support of this hypothesis, the association of the facultative heterochromatin mark H3K27me3 with the HSV-1 genome increases dramatically between 10- and 15-days post-infection *in vivo* (46). The kinetics of H3K27me3 deposition remain to be investigated *in vitro*, but if they are mirrored this could suggest that active chromatinization and reinforcement of silencing continues even after initial shut-down of viral gene expression. In the cellular context, H3K27me3 is linked with the recruitment of canonical polycomb repressor complex 1 (cPRC1), which may reinforce silencing through long-range chromosomal interactions or 3D compaction (50, 104, 105). It is therefore possible that even following H3K27me3 formation on the genome, there are additional layers of protein recruitment that build up over time. In addition, it is also possible that the accumulation of viral non-coding RNAs expressed in latency could impact cellular pathways resulting in decreased signaling to the viral genome for reactivation. The use of the Stayput-GFP model system will permit these different avenues to be explored.

Our model system recapitulates the hallmarks of reactivation Phase I, which has previously been explored in *in vitro*systems using a DNA replication inhibitor. These data are interesting considering the discrepancies in conclusions drawn about reactivation between *in vitro* and *in vivo* modeling. There is evidence *ex vivo* for a Phase I as all classes of viral gene are expressed in a disordered, non-cascade fashion when a combination of explant and nerve-growth factor deprivation are used (106). However, in other models of reactivation *ex vivo*, there is evidence that Phase-I-like gene expression may not occur, especially from studies investigating the requirement for histone demethylase inhibitors. These latter experiments used explant (axotomy) to induce reactivation and found that the earliest induction of lytic gene expression is dependent on lysine 9 demethylase activity (40, 41). In a recent study from our lab, we have found that Phase I reactivation can occur *ex vivo* when axotomy is combined with PI3-kinase inhibition, although with more rapid kinetics than those observed here (107). These discrepancies between may result from the different trigger used to induce reactivation or currently unknown effects of latency established *in vivo*. Importantly, here we have demonstrated that any potential differences between *in vivo* and *in vitro* observations on the mechanisms of reactivation do not result from the use of ACV to establish a quiescent infection. Further work using the Stayput-GFP model system will help elucidate how changes in the host immune response, neuronal subtype, and stimulus used can potentially alter the mechanisms of viral gene expression during reactivation.

Phase I, in addition to occurring synchronously and independently of histone demethylases, also occurs in the absence of viral protein synthesis. In an *in vitro* model employing ACV during latency establishment and stimulus LY294002 during reactivation, it is demonstrated that initial viral transcription occurs before the appearance of viral late protein synthesis and, specifically, independently of viral transactivator VP16 (25, 51). Therefore, cellular host factors must be responsible for instigating the initial reactivation process. Evidence from an *in vitro* model system has demonstrated these events are in fact navigated by cellular proteins JNK and DLK (52). Interestingly, host cell proteins may also be implicated in restricting the full reactivation process, including Gadd45b which appears to antagonize the HSV-1 late expression program to prevent full reactivation (51). Interestingly, Gadd45b mRNA is increased in response to LY294002 only in infected neurons, suggesting perhaps that a viral factor may be mediating the gatekeeping from Phase I to Phase II reactivation.

In our system, Phase I gene expression was dependent on the neuronal regulator of JNK activity, DLK, highlighting the central role of DLK in HSV-1 reactivation. DLK is a cell protein implicated in neuronal stress signaling upstream of cellular protein JNK (108). It has previously been found to be essential for HSV-1 reactivation following PI3-kinase inhibition (52), as well as neuronal hyper-excitability through forskolin (54). However, it has not, until now, been shown to be central to reactivation mediated by heat shock. Although heat shock has been used as a trigger for HSV-1 reactivation (91–98), the downstream molecular events following this stimulus are not well elucidated. Multiple studies have demonstrated that heat shock during reactivation leads to the up-regulation of heat shock proteins, although none of them knowingly relate to DLK. Following hyperthermia-induced reactivation *in vivo*, heat shock protein HSP60 and HSP40 have been demonstrated to be up-regulated (98). Components of the heat shock response pathway have also been demonstrated to be up-regulated by LY294002 treatment in an *in vitro* system (51), including HSP70. In fact, in this same system, treatment with cultures of heat shock factor 1 (HSF-1) activator compound causes robust reactivation. Outside of the virological context, heat shock protein chaperone HSP90 has been shown to bind and maintain DLK stability *in vivo* and it is specifically required for DLK function following axon injury signaling (109). It is a possibility that heat shock in our system is enhancing the function of DLK. Therefore, multiple signals may converge on DLK, which is then able to activate JNK and protein histone phosphorylation and to promote lytic gene expression from the heterochromatin-associated viral genome for reactivation to occur. Indeed, synergy has been demonstrated to enhance DLK activity in neurons (110). This central role for DLK is especially important as it is largely a neuron-specific protein that regulates the response to multiple forms of stress (111) and is therefore a potential target for novel therapeutics that would prevent HSV-1 gene expression and ultimately reactivation.

## Methods

### Primary neuronal cultures

Sympathetic neurons from the superior cervical ganglia (SCG) of post-natal day 0–2 (P0-P2) CD1 Mice (Charles River Laboratories) were dissected as previously described (52). Sensory neurons from the trigeminal ganglia (TG) or adult (P21-24) CD1 Mice (Charles River Laboratories) were dissected as previously described (112). Rodent handling and husbandry were carried out under animal protocols approved by the Animal Care and Use Committee of the University of Virginia (UVA). Ganglia were briefly kept in Leibovitz’s L 15-Glutamine before dissociation in Collagenase Type IV (1 mg/mL) followed by Trypsin (2.5 mg/mL) for 20 min each at 37 □C. Dissociated ganglia were triturated, and approximately 5,000 neurons per well were plated onto rat tail collagen in a 24-well plate. Sympathetic neurons were maintained in CM1 (Neurobasal Medium supplemented with PRIME-XV IS21 Neuronal Supplement (Irvine Scientific), 50 ng/mL Mouse NGF 2.5S, 2 mM L-Glutamine, and Primocin). Aphidicolin (3.3 mg/mL) was added to the CM1 for the first 5 days post-dissection to select against proliferating cells. Sensory neurons were maintained in the same media supplemented with GDNF (50 ng/ml; Peprotech 450-44), and more Aphidicolin (6.6 mg/mL) was used for the first 5 days as more non-neuronal cells tend to be dissected with this neuron type.

### Establishment and reactivation of latent HSV-1 infection in primary neurons

Neonatal SCGs were infected at postnatal days 6-8 with either Us11-GFP, SCgHZ, or Stayput-GFP at an MOI of 7.5 PFU/cell assuming 5,000 cells/well in DPBS +CaCl2 +MgCl2 supplemented with 1% Fetal Bovine Serum, 4.5 g/L glucose, and either with ou without 10 mM Acyclovir (ACV) for 3 hr at 37 C. Post-infection, inoculum was replaced with CM1 containing with or without 50 mM. For infections with ACV, the ACV was was wasded out 5-6 days post-infection. Reactivation was carried out with LY294002 20 μM, forskolin 60 μM (pulsed for 20 hours), and heat shock (3 hours at 43 □C) in BrainPhys (Stem Cell Technologies) supplemented with 2 mM L-Glutamine, 10% Fetal Bovine Serum, Mouse NGF 2.5S (50 ng/mL) and Primocin. Reactivation was quantified by counting number of GFP-positive neurons or performing Reverse Transcription Quantitative PCR (RT-qPCR) for HSV-1 lytic transcripts.

## Acknowledgements

We thank Dr. Ian Mohr at New York University for the gift of the Us11-GFP virus and the pSXZY-GFP-Us11 plasmid. We thank Gary Cohen at the University of Pennsylvania for SCgHZ and Vero F6 cells. This work was supported by National Institutes of Health grants NS105630 (ARC), T32GM008136 (SD), AI130618, AI147163 and AI151358 (ACW), The Owens Family Foundation (ARC), and a German Centre for Infection Research (DZIF) Associate Professorship (DPD).

## Data Availability

All nanopore sequencing datasets associated with this study are available via the European Nucleotide Archive under the accession PRJEB51869.

