## Supplemental material for "DLK-dependent biphasic reactivation of herpes simplex virus latency established in the absence of antivirals"

### Supplemental Materials and Methods

Reagents

Compounds used in the study are as follows: Acycloguanosine, LY 294002, Forskolin, GNE-3511, GSK-J4, OG-L002. Compound concentrations were used based on previously published IC50s and assessed for neuronal toxicity using the cell body and axon health and degeneration index as previously published (1).

Preparation of HSV-1 virus stocks

Stayput-GFP, as well as SCgHZ, is propagated and titrated on previously constructed gH-complementing F6 cells, which contain copies of the gH gene under the control of an HSV-1 gD promoter, as described in (2). Vero F6 cells were maintained in Dulbecco's Modified Eagle's Medium (Gibco) supplemented with 10% FetaPlex (Gemini BioProducts) and 250 μg/mL of G418/Geneticin (Gibco).

HSV-1 stocks of eGFP-Us11 Patton were grown and titrated on Vero cells obtained from the American Type Culture Collection (Manassas, VA). Cells were maintained in Dulbecco's Modified Eagle's Medium (Gibco) supplemented with 10% FetalPlex (Gemini Bio-Products) and 2 mM L-Glutamine. eGFP-Us11 Patton (HSV-1 Patton strain with eGFP reporter protein fused to true late protein Us11 (3) was kindly provided by Dr. Ian Mohr at New York University.

Construction of Stayput-GFP

Stayput-GFP was created by inserting an eUs11-GFP tag into the previously created gH-deficient HSV-1 SCgHZ virus (strain SC16) through co-transfection of SCgHZ viral DNA and pSXZY-eGFP-Us11 plasmid (3) in Vero F6 cells, Transfections were carried on in 6 well plates using 2 μg of viral DNA and 2 μg of pSXZY-eGFP-Us11 and 8ul of jetPRIME® (Polyplus). GFP-positive plaques were subjected to 4 rounds of plaque purification.

Viral genomic DNA sequencing

Genomic DNA was isolated from stocks of SCgHZ and Stayput-GFP using the Qiagen DNeasy Blood and Tissue kit. 250 ng of gDNA was used as input for the SQK-LSK109 genomic DNA by ligation protocol (Oxford Nanopore Technologies Ltd.) and resulting libraries were loaded onto individual R9.4.1 flongles for 20 hours of sequencing. Resulting raw fast5 datasets were basecalled using Guppy v4.2.2 (-f FLO-MIN106 -k SQK-RNA002) with only reads passing filter used for subsequent analyses. *De novo* assembly was performed as follows. First, HSV-1 reads were extracted from the total pool by aligning against the HSV-1 strain SC16 reference genome (MN159383.1) using MiniMap2. HSV-1 reads shorter than 1,000 nt were subsequently filtered out prior to performing de novo assembly with Canu (4). Canu yielded a single circular contig for both SCgHZ and Stayput-GFP which was manually linearized at the canonical HSV-1 start site. Four rounds of genome polishing was performed using racon (5) before a final round of SNP calling was achieved using medaka (https://github.com/nanoporetech/medaka).

Direct RNA sequencing

For both SCgHZ and Stayput-GFP, NHDFs (passage 13) were infected at an MOI of 10. At 6 hours post infection, the supernatant was removed and cells washed once with PBS prior to lysing with 8 ml Trizol. Total RNA was extracted according to Trizol manufacturer's instructions and eluted in nuclease-free water. Nanopore direct RNA-Seq libraries were prepared for each sample using between ~500 ng of poly(A) RNA that was previously isolated from 30 µg of total RNA using the Dynabeads™ mRNA Purification Kit (Invitrogen, 61006). The poly(A) RNA was spiked with 0.5 µL of a synthetic Enolase 2 (ENO2) calibration RNA (Oxford Nanopore Technologies Ltd.) and direct RNA-Seq libraries prepared according to the standard SQK-RNA002 protocol (Oxford Nanopore Technologies Ltd.). Raw fast5 datasets were then basecalled using Guppy v4.2.2 (-f FLO-MIN106 -k SQK-RNA002) with only reads passing filter used for subsequent analyses. Sequence reads were aligned against the Stayput-GFP genome using MiniMap2 (Li, 2018) (-ax splice -k14 -uf --secondary=no), with subsequent parsing through SAMtools and BEDtools (6, 7). Here sequence reads were filtered to retain only primary alignments (Alignment flag 0 (top strand) or 16 (bottom strand)). Figures associated with this study were generated using the R packages Gviz (8) and GenomicRanges (9).

Single-step growth curve

1 x $10^5$ Vero F6 cells were seeded per well in a 24-well plate. 24 hours later, when the cells had reached approximately 90% confluency, cells were infected with Stayput-GFP, SCgHZ, or wild-type Patton Us11-GFP at an MOI of 5 PFU/cell for 1 hour at 37°C with gentle rocking (60 rpm). Inoculum was then removed and replaced with 400 μL of media. At the specified time-points, 400 μL of 9% sterile milk was added to the well and the plates were frozen in the -80C. Following three freeze thaw cycles, titrations were carried out on Vero F6 cells.

Immunofluorescence

Neurons were fixed for 15 min in 4% Formaldehyde and blocked in 5% Bovine Serum Albumin and 0.3% Triton X-100 and incubated overnight in primary antibody. Following primary antibody treatment, neurons were incubated for 1 hr in Alexa Fluor 488-, 555-, and 647-conjugated secondary antibodies for multi-color imaging (Invitrogen). Nuclei were stained with Hoechst 33258 (Life Technologies). Images were acquired using an sCMOS charge-coupled device camera (pco.edge) mounted on a Nikon Eclipse Ti Inverted Epifluorescent microscope using NIS-Elements software (Nikon). Images were analyzed using ImageJ.

Analysis of mRNA expression and viral genome copy number

To assess expression of HSV-1 lytic mRNA, total RNA was extracted from approximately 5,000 neurons using the Quick-RNA Miniprep Kit (Zymo Research) with an on-column DNase I digestion. mRNA was converted to cDNA using the Maxima cDNA synthesis kit (Thermo Fisher) using random hexamers for first-strand synthesis and equal amounts of RNA (20–30 ng/ reaction). To assess viral DNA load, total DNA was extracted from approximately 5,000 neurons using the Quick-DNA Miniprep Plus

Kit (Zymo Research). qPCR was carried out using PowerUp$^{TM}$ SYBR Green Master Mix (Thermoscientific). Relative mRNA/DNA copy numbers were determined using the Comparative CT (ΔΔCT) method normalized to mRNA levels in latently infected samples. Viral RNAs were normalized to mouse reference gene 18s ribosomal RNA. All samples were run in duplicate on an Applied Biosystems QuantStudio 6 Flex Real-Time PCR System and the mean fold change compared to the reference gene calculated. Exact copy numbers were determined by comparison to standard curves of known DNA copy number of viral genomes or plasmid containing 18S rRNA (p18S.1 tag was a gift from Vincent Mauro; Addgene plasmid # 51729 ; http://n2t.net/addgene:51729 ; RRID:Addgene_51729, (10).

Statistical Analysis

All statistical analyses were performed using Prism V8.4. The normality test and statistical test used for each figure are denoted in the figure legends. Individual biological replicates are plotted, each corresponding to a single well of cells/neurons. All neuronal experiments were repeated from pooled neurons from at least 3 litters.

**Supplemental data**

**Supplemental FIG 1**: Stayput-GFP is unable to spread.
Supplemental to Figure 1.

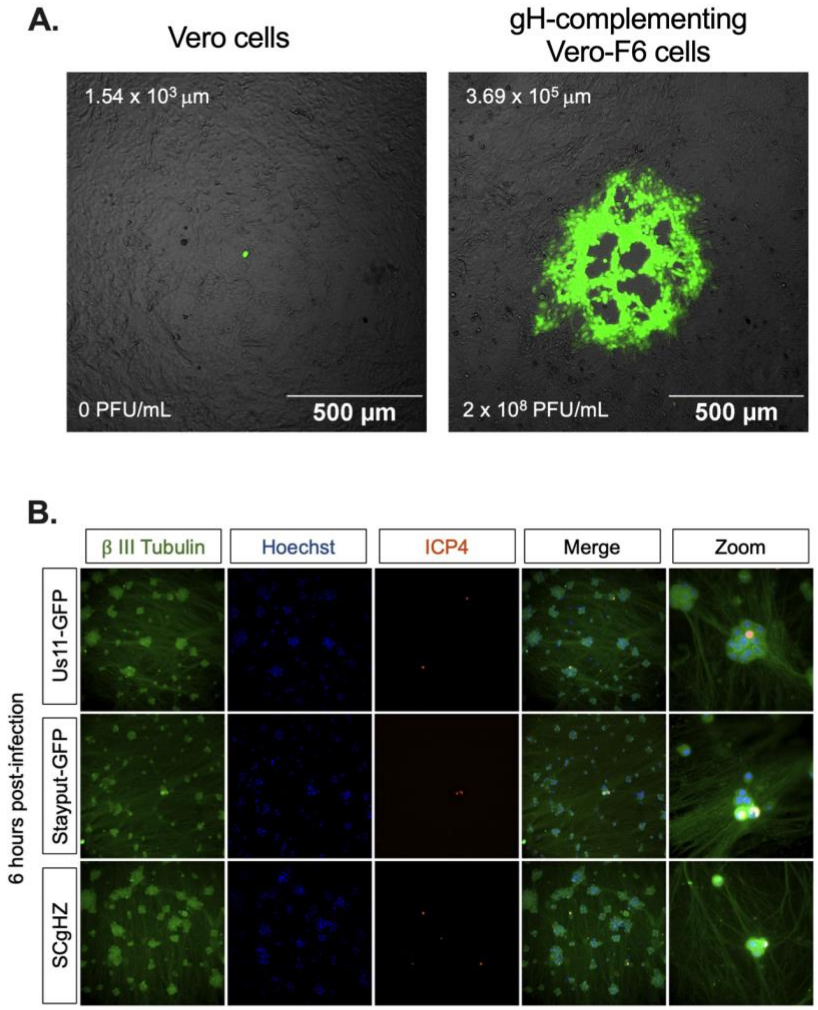


A). Plaque forming assay of Stayput-GFP on Vero (left) or gH-complementing Vero-F6 (right) cells 48 hours post-infection at a $10^{-7}$ dilution. GFP-positive area in example image reported top left of each image. Titer of viral stock reported bottom left of each image.
B) Neonatal sympathetic neurons were infected with Stayput-GFP, SCgHZ, or Us11-GFP at an MOI of 0.5 PFU/cell. Neurons were fixed at 6 hours post-infection and stained for viral protein ICP4 (red) along controls β III tubulin (green) and Hoescht stain (blue).

**Supplemental FIG 2**: Stayput-GFP can be used to create a quiescence model in the absence of viral DNA replication inhibitors in adult sensory neurons.
Supplemental to figure 2.

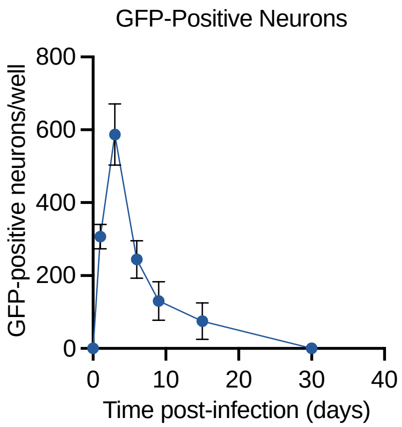


Neurons isolated from the trigeminal ganglia (TG) of female mice were infected with Stayput-GFP at an MOI of 7.5 PFU/cell. The resolution of lytic infection was monitored over time by imaging and counting GFP-positive neurons. n=12, 3 biological replicates. The mean and SEM are shown.

**Supplementary FIG 3**: Viral protein synthesis can be restarted following latency establishment.

Supplemental to figure 4.

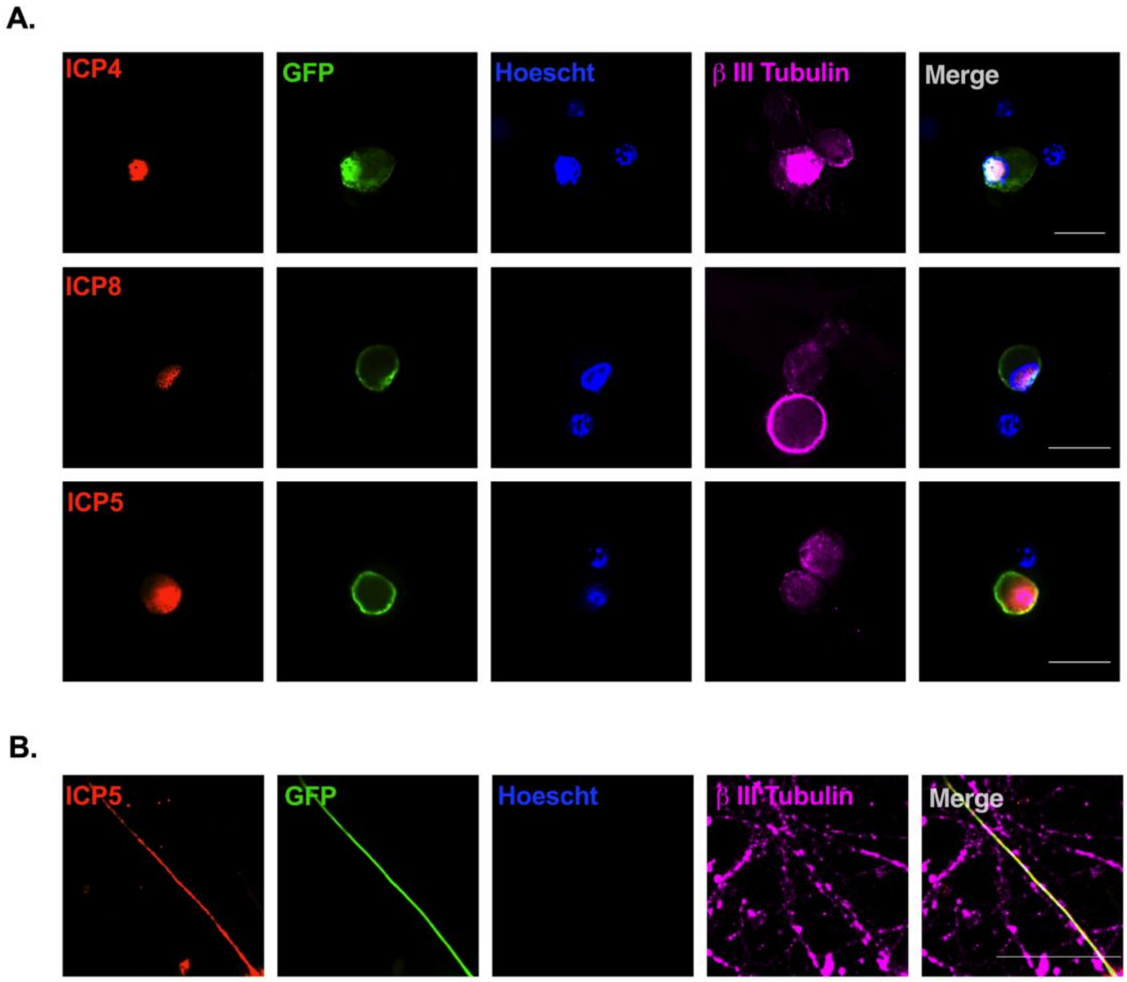


Following infection with Stayput-GFP at MOI 7.5 PFU/cell and the loss of GFP, neonatal SCGs were triggered with LY294002, forskolin, and heat shock. Neurons were fixed at 48 hours post-stimulus and stained with viral immediate early (*ICP4*, A), early (*ICP8*, B), or late (*ICP5*, C-D) protein (red), GFP (green), Hoescht (blue), and β II tubulin (magenta). 25 μm scale bar.

**<u>Supplementary FIG 4:</u>** <u>Additional viral transcripts in neonatal SCG reactivation.</u>
Supplemental to figure 4

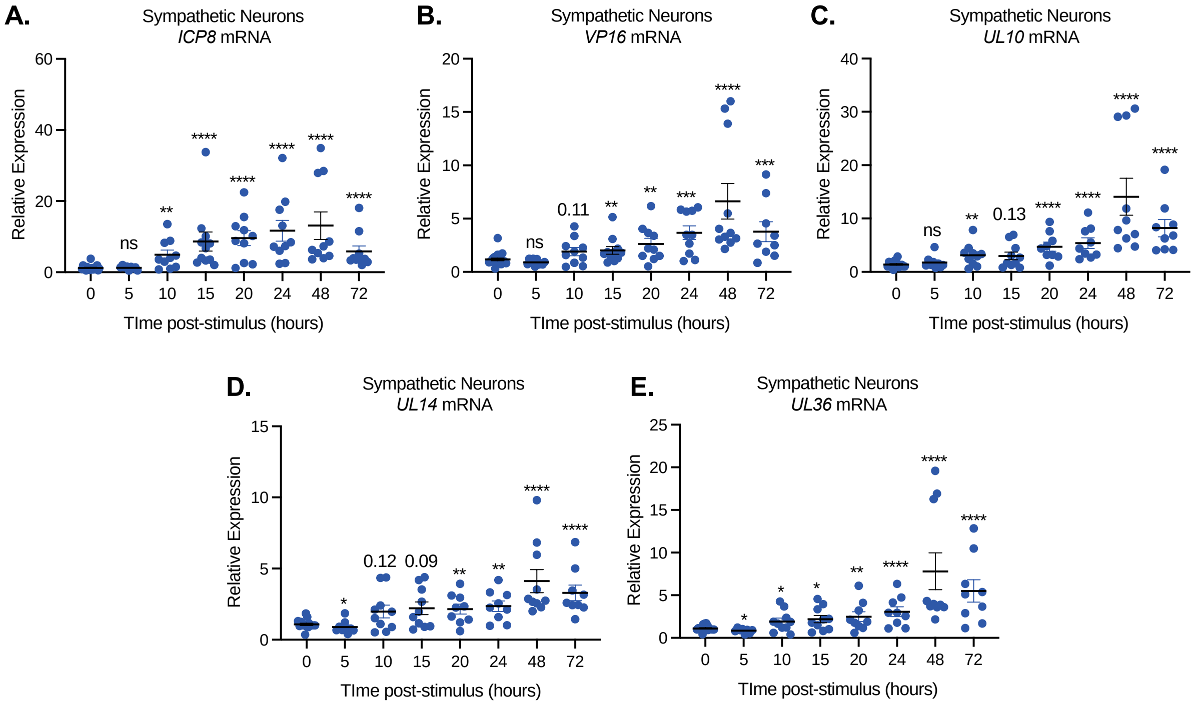


Following infection with Stayput-GFP at MOI 7.5 PFU/cell and the loss of GFP, neonatal SCGs were triggered with LY294002, forskolin, and heat shock. An additional early transcript (A) and late transcripts (B-E) were monitored over time. n=12, 3 biological replicates, Mann-Whitney against Mock. *p<0.05, **p<0.01, ***p<0.001, ****p<0.0001. Individual biological replicates along with the means and SEMs are shown.

**Supplementary FIG 5**: Additional viral transcripts in adult TG reactivation.
Supplemental to figure 4

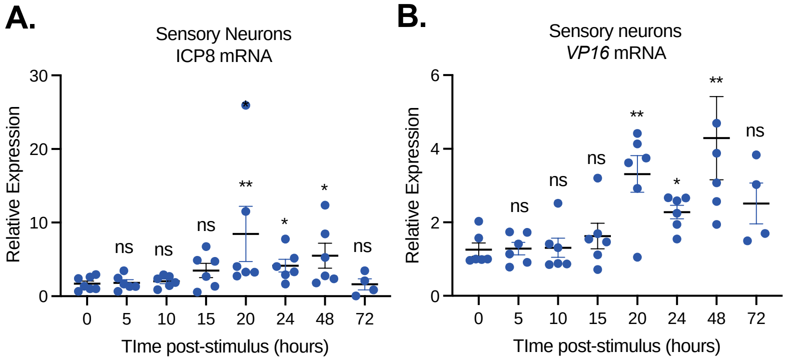


Following infection with Stayput-GFP at MOI 7.5 PFU/cell and the loss of GFP, adult TGs were triggered with LY294002, forskolin, and heat shock. An additional early transcript (A) and late transcript (B) were monitored over time. n=6, 2 biological replicates, Mann-Whitney against Mock. *p<0.05, **p<0.01, ***p<0.001, ****p<0.0001. Individual biological replicates along with the means and SEMs are shown.
